# Signal Peptide of Human Serum Albumin (residues 1-18) Forms Amyloid-like Aggregates

**DOI:** 10.1101/2023.10.24.563726

**Authors:** D. C. Thakur, Kumar Udit Saumya, Vipendra Kumar Singh, Rajanish Giri

## Abstract

Signal peptides (SPs) are essential tools in sorting or translocating the synthesized proteins. However, except few examples, their studies in isolation are quite limited. Here we asked a question of the aggregation propensity of signal peptide (residues 1-18) of Human Serum Albumin (HSA). Molecular dynamic simulations were used to acquire observations into the mechanism through which signal peptides of HSA self-aggregated in physiological conditions. Further, we confirmed our results by combining dye-based assays, atomic force microscopy, and transmission electron microscopy techniques. It was noted that the signal peptide of HSA (region 1-18) forms typical amyloid-like fibrils.

**Graphical:** 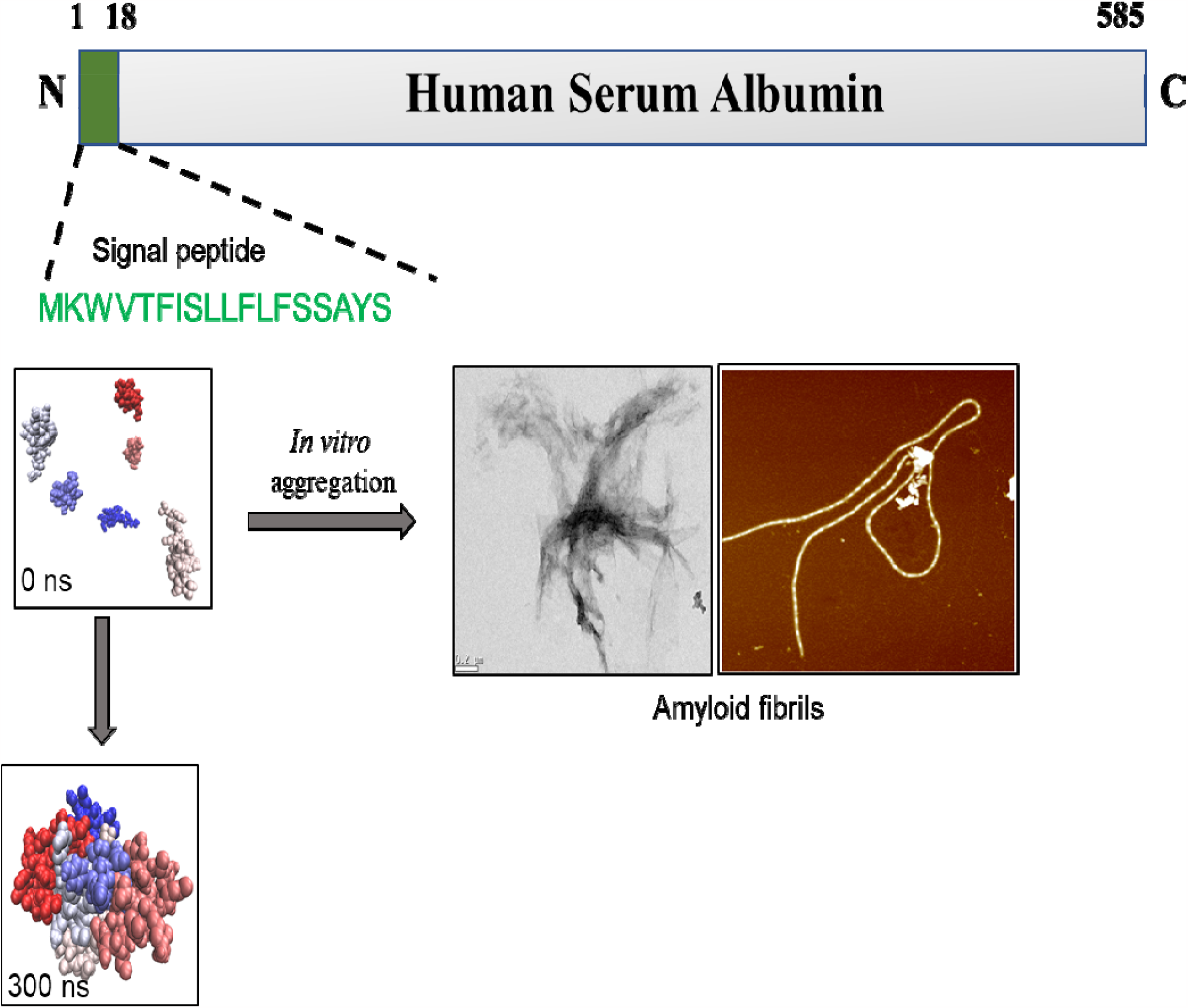

## Introduction

Human serum albumin (HSA), the most abundant protein in the body, performs essential biological functions and binds to various ligands, including amino acids, fatty acids, ions, and hormones (PMID: 26055641; PMID: 30012492; PMID: 25674083). The half-life of HSA is approximately 20 days, and they are present in lymph, saliva, cerebrospinal, intestinal fluid, and plasma (Rabbani & Ahn, 2019; burek et al., 2011). Human serum albumin has worked as a molecular chaperone whose major function is restricting the aggregation of other proteins. They also protect enzymes from thermal degradation and aggregation (PMID: 20133158).

The signal peptides (SPs) are short peptides located at the N-terminal of proteins, involved in protein transport and release (PMID: 35118058). They are present in all eukaryotes and prokaryotes and have diverse applications in various industrial and scientific fields (PMID: 29958716). They are first identified by the signal recognition particle (SRP), a ribonucleoprotein complex, and subsequently enter between the twin transmembrane domain of Sec 61a (PMID: 9823886). In eukaryotes, the process starts when the SRP allows the N-terminal domain of a secretory protein. This episode permits co-translational shift of the protein into the lumen of endoplasmic reticulum (PMID: 11148214). Post transfer of the SPs into the lipid membrane, they are dissected by the membrane-bound SPs and subsequently break down by intermembrane proteases (PMID: 21810987).

The HSA signal peptide (residues 1-18) plays a crucial role in the folding and activity of recombinant proteins folded by 17 disulfide bonds (PMID: 29958716). Signal peptides are important molecular entities with a canonical function in targeted secretion translocation of proteins. They are also significantly involved in the production of recombinant proteins. Most of the aggregation associated studies have been demonstrated the aggregation propensity of HSA protein in the presence of metal ions or alone (PMID: 30384272; PMID: 32023900). However, the misfolding and self-aggregating-like activity of signal peptides of HSA in isolation still pose unanswered questions (PMID: 29958716). Understanding the aggregation potential of signal peptides of HSA may provide valuable insights into the pathophysiology of related diseases. The formation of amyloid, a multi-step process that can lead to the production of highly toxic intermediates in biological systems, is associated with various diseases such as Alzheimer’s, systemic amyloidoses, synucleinopathies, and type 2 diabetes mellitus (PMID: 30384272; PMID: 32023900; PMID: 19049981; PMID: 20079882; https://doi.org/10.1016/j.xcrp.2021.100599). Analyzing the aggregation propensity of the signal peptide of HSA can contribute to a better understanding of disease pathophysiology.

The aggregation of the signal peptide of HSA protein after its delivery to the intended destination raises questions about the consequences if the protein quality control system fails to degrade this cleaved-off signal peptide (PMID: 34050864). We aimed to investigate the fate of the signal peptide if it is not degraded by the protein homeostasis machinery or the protein quality control system and hypothesized that it may aggregate. Due to its solubility only in DMSO, studying the monomer was challenging. To test this hypothesis, we used in-silico tools, MD simulations, dye-based spectroscopy, and microscopy to predict and confirm the self-aggregation propensity of the HSA signal peptide (region 1-18) under physiological conditions.

## 2 Materials and Methods

### 2.1 Reagents

Signal peptide (residues 1-18) “MKWVTFISLLFLFSSAYS” of HSA protein was obtained from NCBI, chemically synthesized, and purchased from Genscript, USA, with 91.5% purity. TEM grid was acquired from TED PELLA INC., USA. Thioflavin T (ThT), Congo Red (CR), 8-Anilinonaphthalene-1-sulfonic acid (ANS), 1,1,1,3,3,3-hexafluoro-2-propanol (HFIP), were acquired from Sigma-Aldrich (St. Louis, MO, USA). Other chemicals used in this study were of the highest purity acquired from regional sources.

### 2.2 In silico prediction of fibril formation

The aggregation score (aggregation propensity region) of the signal peptide of HSA protein was predicted using different freely accessible data servers such as FoldAmyloid, Metamyl, TANGO, AGGRESCAN, and CamSol (Fernandez-Escamilla et al., 2004; Pawlicki et al., 2008; Conchillo-Solé et al., 2007; Sormanni et al., 2015; Garbuzynskiy et al., 2010). In short, parameters such as physiological setup (Temp-37°C and pH-7.4) were used to run the TANGO software. Whereas other databases, namely CamSol, Aggrescan Foldamyloid, and NHSA, were used in default settings. We used Peptide 2.0 (www.peptide2.com) to predict hydrophobicity, and the physiological parameters of the signal peptide were calculated with the help of the PROTPARAM database (www.expasy.org/protparam).

#### MD Simulation of HSA signal peptide (residues 1-18)

The simulations utilized the GROMACS software package (version 2022.3) along with the GROMOS96 54a7 force field (Schmid et al., 2019). The SPC (simple point charge) water model was used, and periodic boundary conditions (PBC) were implemented using a cubic simulation box. To ensure a minimum initial distance of 1 nm between proteins and the box boundaries, the system was set up accordingly. Nonbonded interactions were truncated at a 1.2 nm cutoff, and long-range electrostatics were computed using the Particle-mesh Ewald (PME) summation method [DOI: 10.1021/acs.jpcb.5b04822]. The temperature was maintained at 300 K using a modified Berendsen thermostat (V-rescale), while the pressure was regulated at 1 bar with the Parrinello-Rahman barostat (Ng et al., 2021). Counterions were added to neutralize the system, and additional salt was included to achieve the desired concentration. Prior to the main simulations, energy minimization employing the steepest descent algorithm was conducted, followed by equilibration stages consisting of a 10 ns NVT simulation and a subsequent 10 ns NPT simulation. During equilibration, the protein backbone was restrained to maintain stability. These preliminary simulations were performed to generate the initial configurations necessary for the aggregation simulations. The starting conformation for the aggregation simulations was obtained from the 1 microsecond monomer simulations of HSA_SP (1-18). In aggregation simulation, six monomers were randomly placed in a periodic cubic box with an edge length of 10 nm, corresponding to a concentration of 0.15 M. The systems were solvated, and appropriate ions were added to neutralize the system and achieve the desired salt concentration. Before the production run, the aggregation simulations were further equilibrated using a 30 ns NVT simulation, followed by a 30 ns NPT simulation. The aggregation simulation was run for 300 ns After the equilibration (Samantray et al., 2022; Carballo-Pacheco, and Strodel, 2016).

### 2.3 Thioflavin T assay

In vitro, aggregation was estimated using fluorescent dye-based assays such as ANS and ThT. A 200 μM ThT stock solution was prepared in distilled water, while a final concentration of 20 μM was used for a dye-based binding assay. Data was acquired using a black 96-well flat-bottomed plate through a microplate reader (TECAN Infinite M200 PRO). The excitation wavelength of 450 nm and emission spectra from 470 nm to 700 nm was used. Experiments were performed at least in triplicates.

### 2.4 ANS-dye-based fluorescence assay

ANS-dye-based binding analysis of signal peptide of HSA protein was conducted using a 20 μM concentration. A final concentration of 30 μM for both monomer and aggregated signal peptide of HSA protein was used in sodium phosphate buffer (20 mM; pH 7.4). Data were acquired in the TECAN Infinite M200 PRO microplate reader. The excitation wavelength of 380 nm and emission spectra ranging between 400 to 650 nm were used. The control used in the experiment was ANS dye only with buffer. Experiments were performed at least in triplicates.

### 2.5 Congo Red shift assay

Congo red (CR) dye is amyloid specific that binds with cross β-sheet rich amyloid-like aggregates and shows characteristic redshift in spectra. Mili-Q water was used to prepare a stock solution (0.1% w/v) of CR. Then CR solution with a concentration of 25 mM was added to the aggregated sample of the signal peptide of HSA protein (residues 1-18). Tecan Infinite M200 microplate reader was used to record the absorption spectra at 400-450 nm wavelength using a transparent 96-well plate. Experiments were performed at least in triplicates

### 2.6 High-Resolution Transmission Electron Microscopy (HRTEM).

For TEM analysis, 7 μL of the aggregate sample (diluted 10× in Milli-Q) was placed on 200 mesh grids of copper and incubated at RT for 60 s. The grid was rinsed softly, swirling in water droplets for at least 30 s three times afterward. An aqueous solution of ammonium molybdate (3% w/v) was used to stain the grids negatively. In the end, the grids were allowed to dry overnight, and then a snapshot was captured in HRTEM (FP 5022/22-Tecnai G2 20 S-TWIN, FEI) at 200kV accelerating voltage.

### 2.7 Atomic Force Microscopy (AFM).

The aggregate sample was first diluted 30 x times with Milli-Q and then drop casted using 7 μL on a newly dissected mica sheet. After 2 min incubation, the specimen was rinsed with deionized water and dried overnight in a desiccator. In the end, a snapshot was captured on AFM at the tapping mode (Dimension Icon from Bruker).

## 3. Results and Discussion

### 3.1 Prediction of aggregation propensity of the signal peptide (region 1-18) of HSA protein.

The aggregation tendency of every protein is modelled by evolution according to its actual presence in the organisms. HSA is an abundant protein in the human body, and it displays an excellent ligand-binding potential and is also a carrier of various exogenous and endogenous proteins. Protein aggregation is affected by several factors, such as the protein structure (primary, secondary, or tertiary), processing, and the environment of the protein in which protein exists. Aggregation or misfolding may reduce or entirely alter their biological characteristics. Therefore, protein aggregation remains a hurdle in the commercialization and development of biotechnology products. Various in-silico-based platforms that predict APR from the polypeptide sequence of signal peptides of HSA based on biochemical and physicochemical characteristics have been available. In the present study, we have used five web freely available high-precision server tools for APR prediction in the signal peptide of HSA (TANGO, AGGRESCAN, FoldAmyloid, CamSol, and AGGRESCAN). Our in silico server-based results revealed greater prediction scores for aggregation propensity of signal peptide region 1-18, especially the residue between the 2^nd^-15^th^ (Figure 1 A-E). Human serum albumin involves transporting various exogenous and endogenous chemicals such as metals, drugs, amino acids, bile acids, toxic metabolites, hormones, and fatty acids (Taguchi et al., 2012; Otagiri & Chuang, 2009). Signal peptides are key tools that control the function and folding of recombinant proteins, which are folded by disulfide bonds. Human serum albumin is a multifunctional protein with wide application in clinical settings.

**Figure 1:**
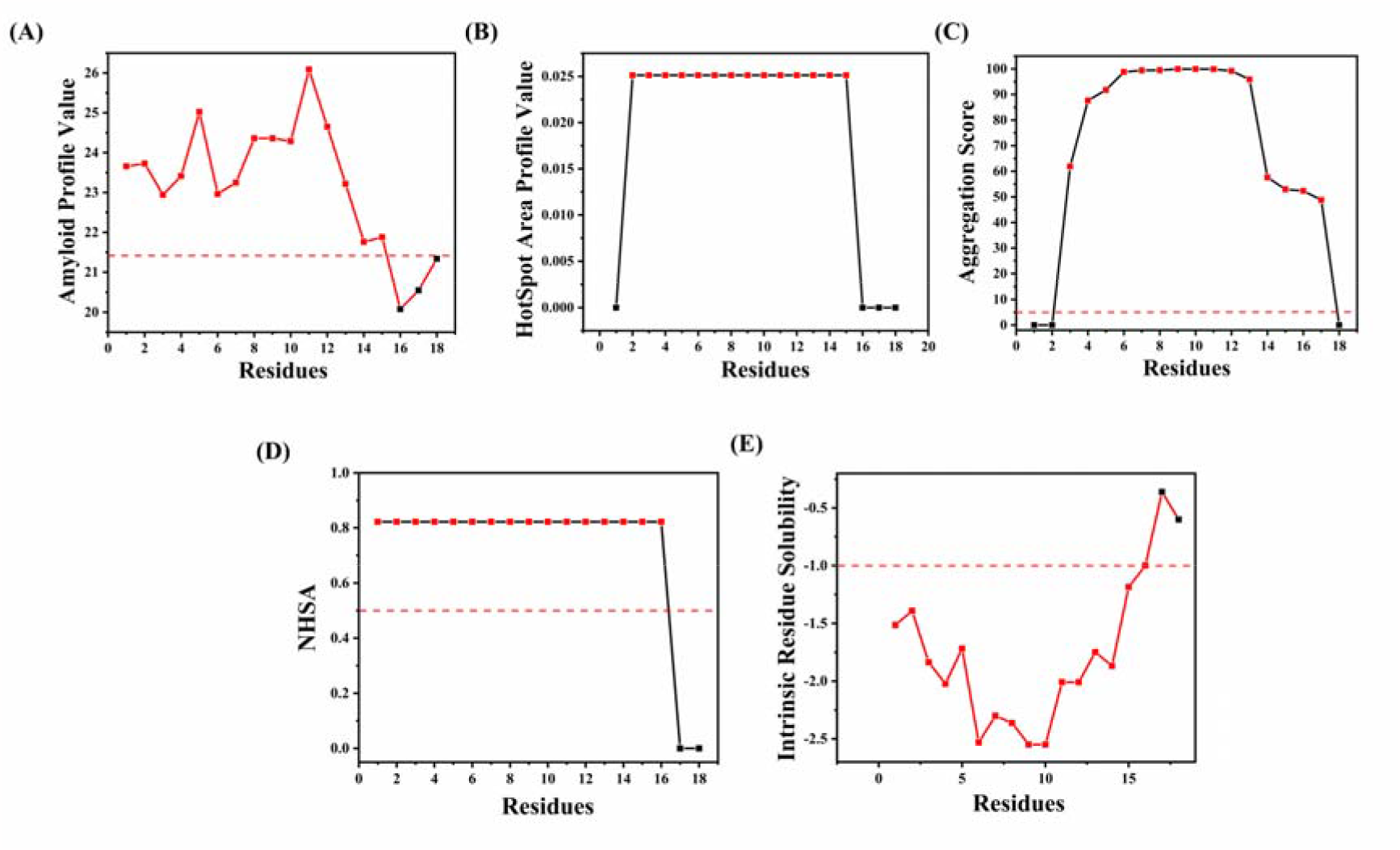
Prediction of aggregation-rich regions in the signal peptide of human serum albumin. Five web server tools predicted APRs of Human serum albumin’s signal peptide sequence (residues 1-18). (A) FoldAmyloid aggregation scan showing information on amyloidogenic region (B) MetAmyl server showing aggregation propensity as hotspot area profile value (C) TANGO server showing the percentage probability of the aggregation-prone region in the signal peptide of HSA. (D) Aggrescan servers showed the normalized hotspot area for APRs (E) CamSol solubility scan showed the solubility of the signal peptide of HSA.

The physicochemical properties of our signal peptide sequence were investigated utilizing Protparam and www.peptide2.com for the hydrophobicity index, respectively. We observed mostly hydrophobic amino acids, with an overall 11 of them allocated across the whole signal peptide of the HSA sequence (hydrophobicity index: 61.11%) (as shown in Figure 1).

### 3.2 Prediction of the aggregation of the signal peptide of HSA at the atomistic scale

Protein aggregation of signal peptides of HSA is a discussion of great concern to the researcher due to its involvement in protein homeostasis system and protein translocation. Numerous *in-silico* tools are available to predict aggregation propensity from analysis of amino-acid residues in a polypeptide chain. Similarly, investigation of the structural rearrangements during the aggregation and disaggregation process through numerous MD simulation techniques have been reported earlier. In this study, we therefore performed MD simulation of HSA signal peptide (region 1-18) to elucidate its aggregation behavior at the atomic level in presence of multiple monomeric subunits. GROMOS96 54a7 force field was applied to record the aggregation of signal peptides of HSA under physiological temperature. MD simulation was run for 300 ns, and a snapshot of signal peptides is illustrated in Figure 2. After the run of 50 ns, the signal peptides converted into dimers with following trimer, tetramer, pentamer, and at the last converted to hexamer (at 300 ns). So far, any study related to MD simulation that showed the propensity of self-aggregation behavior of signal peptides of HSA (region 1-18) under physiological conditions in not reported to date.

**Figure 2:**
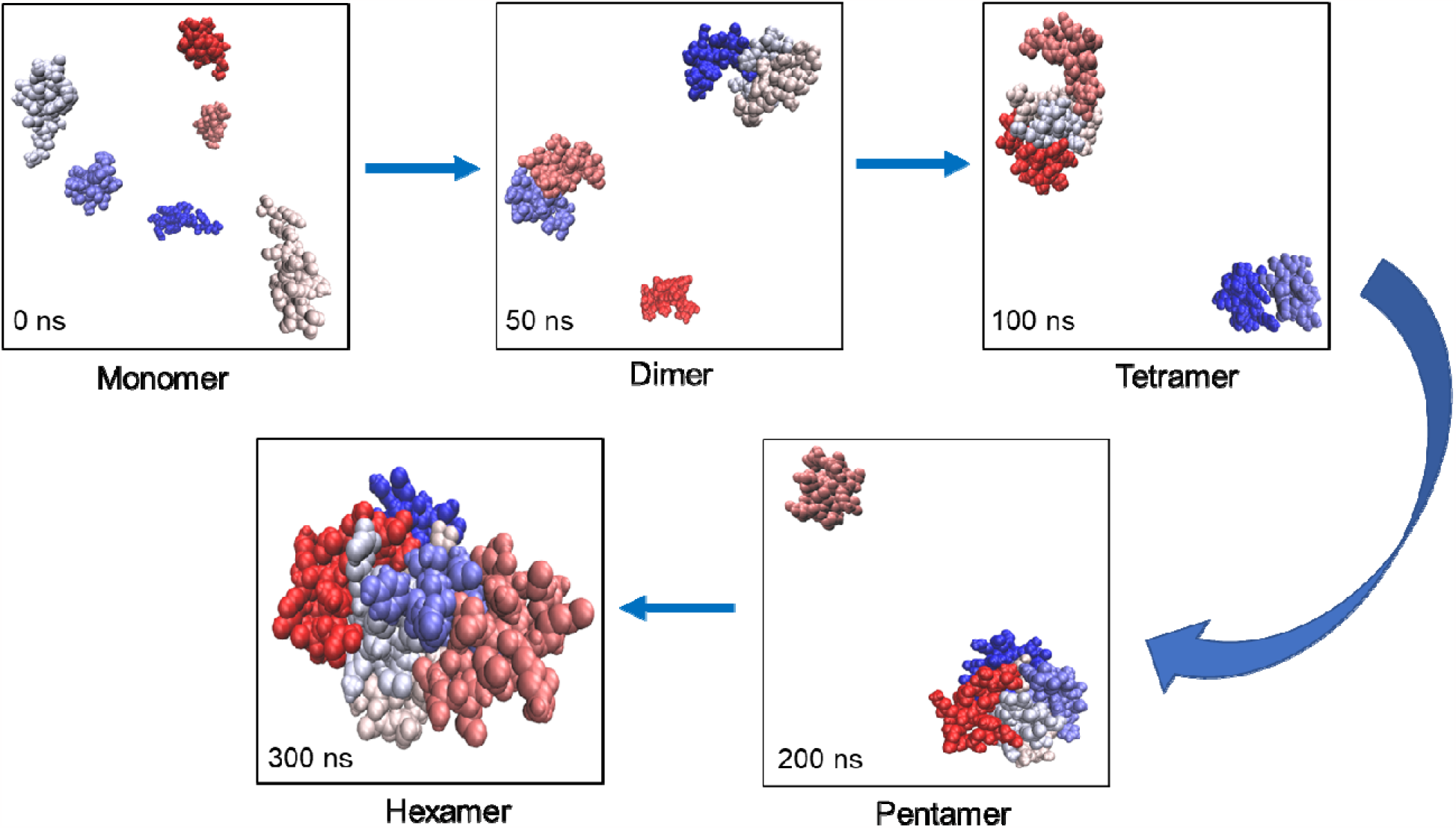
MD simulation analysis of signal peptide of HSA (region 1-18), at atomistic scale snapshot showing the aggregation behaviour from monomer to subsequently dimer, tetramer, pentamer, and hexamers of peptides in functions of time.

### 3.3 Binding of signal peptide sequence of HSA (residues 1-18) to amyloid-specific dyes and enhanced fluorescence.

Fluorescence is one of the most useful techniques for observing and structurally depicting amyloid fibrils or protein aggregation in pathogenic or clinical settings (Gasteiger et al., 2005; Hawe et al., 2008). The most common methods for quantifying and spotting amyloid-like fibrils depend on observing the fluorescence alterations in thioflavin T and 8-Anilinonaphthalene-1-sulfonic acid (ANS) dye and alteration in absorbance of the Congo red (CR), an azo dye. The ANS dye binds to the hydrophobic regions, enhancing fluorescence levels with a typical shift in emission spectra like a blue shift. First, we analyzed the fibril formation of the signal peptide of HSA protein by ANS-dye-based assay (as shown in Figure 2A). The reading was obtained at the initial (0 h) and final (168 h) of the aggregate samples, demonstrating an increased fluorescence intensity of ANS approximately 1.5-fold. At 168 hrs, we observed a significant gain in fluorescence accompanied by a characteristic blue shift (549 nm to 516 nm).

In our study, we found that the signal peptide of the HSA protein shows a significant change in fluorescence signal when compared to monomer and ThT-only control. The aggregates of the signal peptide of HSA demonstrated a ~8-fold increment in ThT fluorescence intensity. It indicates the generation of amyloid-like fibrils in the aggregate-inducing environment under physiological settings, as shown in Figure (2B). We also noticed that CR binds to signal peptides of HSA proteins and demonstrates a significant shift in spectra from 488 to 491 nm (as shown in Figure 2C). The signal peptide of HSA (residues 1-18) was also positive for ANS assay, where we observed an increase in fluorescence spectra accompanied by characteristics of blue-shift in spectra from 541 nm - 498 nm.

**Figure 2:**
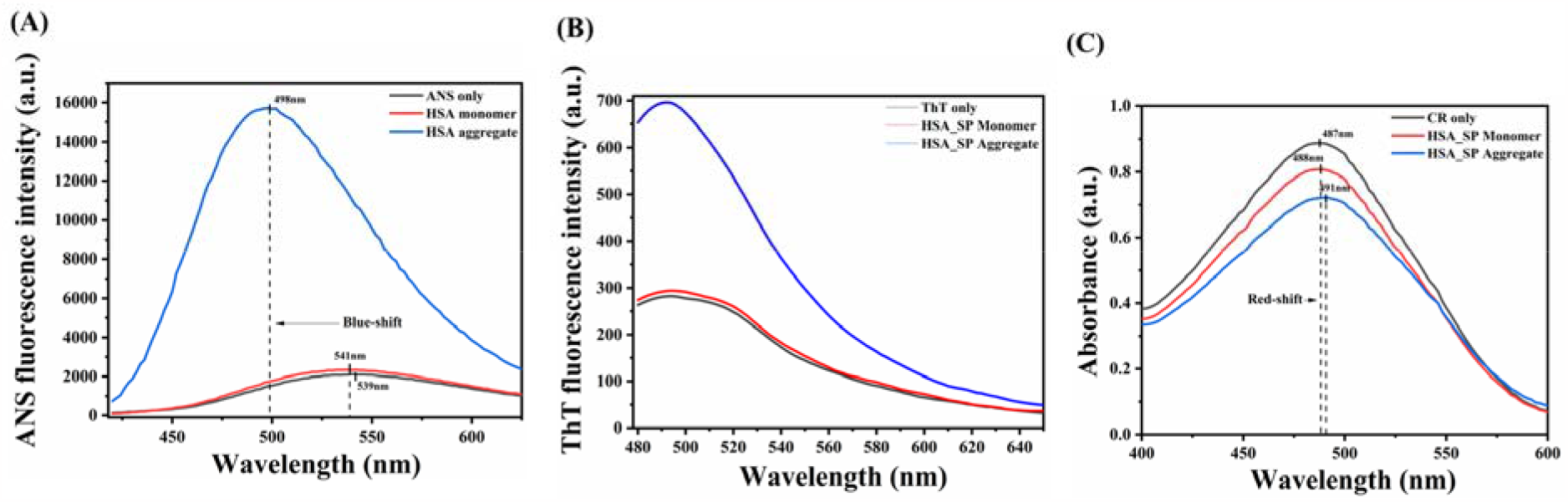
Fluorescent dye binding assay for the analysis of signal peptides of HSA. (A) ANS fluorescence spectra of aggregated signal peptide sample, SP monomer, and ANS dye alone, respectively. The changes in spectra of the monomeric and aggregated samples are shown using an arrow and flecked line (B) ThT fluorescence spectra of the HSA (C) Congo Red signal peptide spectra of the aggregated signal peptide of HSA.

### 3.4 Signal peptides of HSA (residues 1-18) display fibril-like structure in morphology assessment of AFM and TEM.

Atomic force microscopy and High Resolution-TEM are versatile techniques for analyzing protein aggregation in vitro environments (Charnley et al., 2018; Sung et al., 2015). In our study, we analyzed and quantitatively computed the height of amyloid fibrils of our signal peptide samples (as shown in Figure 3A). The height of the fibrils of the signal peptide was approximately 20 nm [Figure 3A(III)]. Next, we confirmed the amyloid fibril formation of our aggregated sample by using transmission electron microscopy. A snapshot of the TEM image clearly showed a distinctive amyloid-like fibrillar structure with protruding short and long dense overlapped fibrils (as shown in Figure 3B).

**Figure 3:**
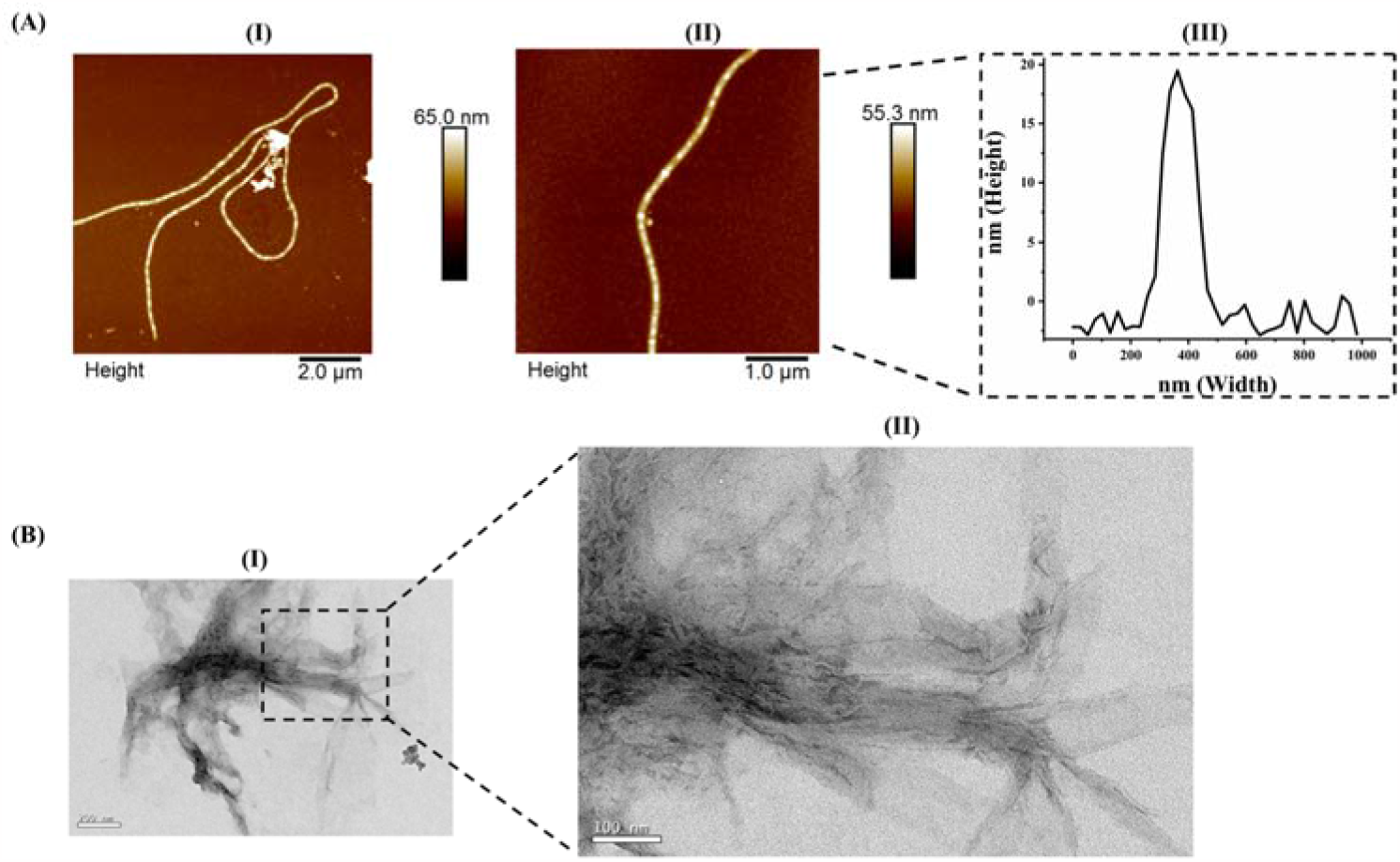
Morphology analysis of signal peptide of human serum albumin amyloid fibrils. (A) AFM analysis represented the height (Y-axis) and width (X-axis) of aggregated signal peptide samples of HSA, where the mesh denoted the fibril used for measuring height. (B) HR-TEM snapshot image captured at 200 kV accelerating voltage.

## 4. Conclusions

From our initial prediction, we found the first 18 residues of HSA to have high aggregation propensity. We utilized the MD simulation to understand the self-aggregation nature of signal peptides of HSA (residues). Our MD simulation study clearly demonstrated the aggregation nature of signal peptides of HSA in the physiological environment. Further, our morphological analysis clearly verified the formation of typical amyloid-like fibrils. In the near future, it will help to understand the mechanisms of the entire HSA protein and its involvement in the transportation of various endogenous and exogenous substances. Our findings also indicate a possible role towards imbalance of HSA signal peptide in amyloidogenic diseases.

## Credit authorship contribution statement

**D.C. Thakur, Kumar Udit Saumya, Vipendra Kumar Singh:** Data acquisition, analysis and interpretation of results, and manuscript writing. **Rajanish Giri:** Conceptualization, Project administration, Supervision, Validation, review & editing of the final manuscript.

## Acknowledgment

D.C. Thakur acknowledges the Indian Council of Medical Research Fellowship, New Delhi, India, and Kumar Udit Saumya acknowledges the Indian Institute of Technology Mandi for Ph.D. Fellowship. The authors thank the Indian Institute of Technology Mandi (Bio X, AMRC, and C4DFED Center) for the facilities. The authors highly acknowledge the National Supercomputing Mission (NSM) for providing computing resources of ‘PARAM Himalaya’ at IIT Mandi, which is implemented by C-DAC and supported by the Ministry of Electronics and Information Technology (MeitY) and Department of Science and Technology (DST), Government of India.

## References

(1) Fanali, G.; di Masi, A.; Trezza, V.; Marino, M.; Fasano, M.; Ascenzi, P. Human Serum Albumin: From Bench to Bedside. Mol Aspects Med 2012, 33 (3), 209–290. 10.1016/j.mam.2011.12.002.

(2) Rabbani, G.; Ahn, S. N. Structure, Enzymatic Activities, Glycation and Therapeutic Potential of Human Serum Albumin: A Natural Cargo. Int J Biol Macromol 2019, 123, 979–990. 10.1016/j.ijbiomac.2018.11.053.

(3) Gburek, J.; Gołąb, K.; Juszczynska, K. [Renal Catabolism of Albumin - Current Views and Controversies]. Postepy Hig Med Dosw (Online) 2011, 65, 668–677. 10.5604/17322693.964329.

(4) Maciążek-Jurczyk, M.; Janas, K.; Pożycka, J.; Szkudlarek, A.; Rogóż, W.; Owczarzy, A.; Kulig, K. Human Serum Albumin Aggregation/Fibrillation and Its Abilities to Drugs Binding. Molecules 2020, 25 (3). 10.3390/molecules25030618.

(5) Mishra, V.; Heath, R. J. Structural and Biochemical Features of Human Serum Albumin Essential for Eukaryotic Cell Culture. Int J Mol Sci 2021, 22 (16). 10.3390/ijms22168411.

(6) Dockal, M.; Carter, D. C.; Rüker, F. Conformational Transitions of the Three Recombinant Domains of Human Serum Albumin Depending on PH. J Biol Chem 2000, 275 (5), 3042–3050. 10.1074/jbc.275.5.3042.

(7) Yang, F.; Phillips, G. N. Crystal Structures of CO-, Deoxy- and Met-Myoglobins at Various PH Values. J Mol Biol 1996, 256 (4), 762–774. 10.1006/jmbi.1996.0123.

(8) Basova, L. v; Il’ina, N. B.; Vasilenko, K. S.; Tiktopulo, E. I.; Bychkova, V. E. [Conformational States of the Water-Soluble Fragment of Cytochrome B5. I. PH-Induced Denaturation]. Molekuliarnaia biologiia 2002, 36 (5), 891–900.

(9) Finn, T. E.; Nunez, A. C.; Sunde, M.; Easterbrook-Smith, S. B. Serum Albumin Prevents Protein Aggregation and Amyloid Formation and Retains Chaperone-like Activity in the Presence of Physiological Ligands. J Biol Chem 2012, 287 (25), 21530–21540. 10.1074/jbc.M112.372961.

(10) de Simone, G.; di Masi, A.; Ascenzi, P. Serum Albumin: A Multifaced Enzyme. Int J Mol Sci 2021, 22 (18). 10.3390/ijms221810086.

(11) Marini, I.; Moschini, R.; del Corso, A.; Mura, U. Chaperone-like Features of Bovine Serum Albumin: A Comparison with Alpha-Crystallin. Cell Mol Life Sci 2005, 62 (24), 3092–3099. 10.1007/s00018-005-5397-4.

(12) Rambaran, R. N.; Serpell, L. C. Amyloid Fibrils: Abnormal Protein Assembly. Prion 2008, 2 (3), 112–117. 10.4161/pri.2.3.7488.

(13) Fassler, J. S.; Skuodas, S.; Weeks, D. L.; Phillips, B. T. Protein Aggregation and Disaggregation in Cells and Development. J Mol Biol 2021, 433 (21), 167215. 10.1016/j.jmb.2021.167215.

(14) Bucciantini, M.; Giannoni, E.; Chiti, F.; Baroni, F.; Formigli, L.; Zurdo, J.; Taddei, N.; Ramponi, G.; Dobson, C. M.; Stefani, M. Inherent Toxicity of Aggregates Implies a Common Mechanism for Protein Misfolding Diseases. Nature 2002, 416 (6880), 507–511. 10.1038/416507a.

(15) Ke, P. C.; Zhou, R.; Serpell, L. C.; Riek, R.; Knowles, T. P. J.; Lashuel, H. A.; Gazit, E.; Hamley, I. W.; Davis, T. P.; Fändrich, M.; Otzen, D. E.; Chapman, M. R.; Dobson, C. M.; Eisenberg, D. S.; Mezzenga, R. Half a Century of Amyloids: Past, Present, and Future. Chem Soc Rev 2020, 49 (15), 5473–5509. 10.1039/c9cs00199a.

(16) Chiti, F.; Dobson, C. M. Protein Misfolding, Functional Amyloid, and Human Disease. Annu Rev Biochem 2006, 75, 333–366. 10.1146/annurev.biochem.75.101304.123901.

(17) Chiti, F.; Dobson, C. M. Protein Misfolding, Amyloid Formation, and Human Disease: A Summary of Progress Over the Last Decade. Annu Rev Biochem 2017, 86, 27–68. 10.1146/annurev-biochem-061516-045115.

(18) Zhou, B.-R.; Liang, Y.; Du, F.; Zhou, Z.; Chen, J. Mixed Macromolecular Crowding Accelerates the Oxidative Refolding of Reduced, Denatured Lysozyme: Implications for Protein Folding in Intracellular Environments. J Biol Chem 2004, 279 (53), 55109–55116. 10.1074/jbc.M409086200.

(19) van den Berg, B.; Ellis, R. J.; Dobson, C. M. Effects of Macromolecular Crowding on Protein Folding and Aggregation. EMBO J 1999, 18 (24), 6927–6933. 10.1093/emboj/18.24.6927.

(20) Munishkina, L. A.; Cooper, E. M.; Uversky, V. N.; Fink, A. L. The Effect of Macromolecular Crowding on Protein Aggregation and Amyloid Fibril Formation. J Mol Recognit 2004, 17 (5), 456–464. 10.1002/jmr.699.

(21) Nguyen PH, Ramamoorthy A, Sahoo BR, et al. Amyloid Oligomers: A Joint Experimental/Computational Perspective on Alzheimer’s Disease, Parkinson’s Disease, Type II Diabetes, and Amyotrophic Lateral Sclerosis. Chem Rev. 2021;121(4):2545–2647. doi:10.1021/acs.chemrev.0c01122

(22) Carballo-Pacheco M, Strodel B. Advances in the Simulation of Protein Aggregation at the Atomistic Scale. J Phys Chem B. 2016;120(12):2991–2999. doi:10.1021/acs.jpcb.6b00059

(23) Ilie IM, Caflisch A. Simulation Studies of Amyloidogenic Polypeptides and Their Aggregates. Chem Rev. 2019;119(12):6956–6993. doi:10.1021/acs.chemrev.8b00731

(24) Sangwan S, Zhao A, Adams KL, et al. Atomic structure of a toxic, oligomeric segment of SOD1 linked to amyotrophic lateral sclerosis (ALS). Proc Natl Acad Sci U S A. 2017;114(33):8770–8775. doi:10.1073/pnas.1705091114

(25) Wong EK, Prytkova V, Freites JA, Butts CT, Tobias DJ. Molecular Mechanism of Aggregation of the Cataract-Related γD-Crystallin W42R Variant from Multiscale Atomistic Simulations. Biochemistry. 2019;58(35):3691–3699. doi:10.1021/acs.biochem.9b00208.

(26) Fernandez-Escamilla, A.-M.; Rousseau, F.; Schymkowitz, J.; Serrano, L. Prediction of Sequence-Dependent and Mutational Effects on the Aggregation of Peptides and Proteins. Nat Biotechnol 2004, 22 (10), 1302–1306. 10.1038/nbt1012.

(27) Pawlicki, S.; le Béchec, A.; Delamarche, C. AMYPdb: A Database Dedicated to Amyloid Precursor Proteins. BMC Bioinformatics 2008, 9, 273. 10.1186/1471-2105-9-273.

(28) Conchillo-Solé, O.; de Groot, N. S.; Avilés, F. X.; Vendrell, J.; Daura, X.; Ventura, S. AGGRESCAN: A Server for the Prediction and Evaluation of “Hot Spots” of Aggregation in Polypeptides. BMC Bioinformatics 2007, 8, 65. 10.1186/1471-2105-8-65.

(29) Sormanni, P.; Aprile, F. A.; Vendruscolo, M. The CamSol Method of Rational Design of Protein Mutants with Enhanced Solubility. J Mol Biol 2015, 427 (2), 478–490. 10.1016/j.jmb.2014.09.026.

(30) Garbuzynskiy, S. O.; Lobanov, M. Y.; Galzitskaya, O. v. FoldAmyloid: A Method of Prediction of Amyloidogenic Regions from Protein Sequence. Bioinformatics 2010, 26 (3), 326–332. 10.1093/bioinformatics/btp691.

(31) Schmid N, Eichenberger AP, Choutko A, et al. Definition and testing of the GROMOS force-field versions 54A7 and 54B7. Eur Biophys J. 2011;40(7):843–856. doi:10.1007/s00249-011-0700-9

(32) Samantray S, Schumann W, Illig AM, et al. Molecular Dynamics Simulations of Protein Aggregation: Protocols for Simulation Setup and Analysis with Markov State Models and Transition Networks. Methods Mol Biol. 2022;2340:235–279. doi:10.1007/978-1-0716-1546-1_12

(33) Carballo-Pacheco M, Strodel B. Advances in the Simulation of Protein Aggregation at the Atomistic Scale. J Phys Chem B. 2016;120(12):2991–2999. doi:10.1021/acs.jpcb.6b00059

(34) Schmid, N.; Eichenberger, A. P.; Choutko, A.; Riniker, S.; Winger, M.; Mark, A. E.; van Gunsteren, W. F. Definition and Testing of the GROMOS Force-Field Versions 54A7 and 54B7. Eur Biophys J 2011, 40 (7), 843–856. 10.1007/s00249-011-0700-9.

(35) Taguchi, K.; Giam Chuang, V. T.; Maruyama, T.; Otagiri, M. Pharmaceutical Aspects of the Recombinant Human Serum Albumin Dimer: Structural Characteristics, Biological Properties, and Medical Applications. J Pharm Sci 2012, 101 (9), 3033–3046. 10.1002/jps.23181.

(36) Otagiri, M.; Chuang, V. T. G. Pharmaceutically Important Pre- and Posttranslational Modifications on Human Serum Albumin. Biol Pharm Bull 2009, 32 (4), 527–534. 10.1248/bpb.32.527.

(37) Gasteiger, E.; Hoogland, C.; Gattiker, A.; Duvaud, S.; Wilkins, M. R.; Appel, R. D.; Bairoch, A. Protein Identification and Analysis Tools on the ExPASy Server. In The Proteomics Protocols Handbook; Humana Press: Totowa, NJ, 2005; pp 571–607. 10.1385/1-59259-890-0:571.

(38) Straub, J. E.; Thirumalai, D. Principles Governing Oligomer Formation in Amyloidogenic Peptides. Curr Opin Struct Biol 2010, 20 (2), 187–195. 10.1016/j.sbi.2009.12.017.

(39) Hawe, A.; Sutter, M.; Jiskoot, W. Extrinsic Fluorescent Dyes as Tools for Protein Characterization. Pharm Res 2008, 25 (7), 1487–1499. 10.1007/s11095-007-9516-9.

(40) Girych, M.; Gorbenko, G.; Maliyov, I.; Trusova, V.; Mizuguchi, C.; Saito, H.; Kinnunen, P. Combined Thioflavin T-Congo Red Fluorescence Assay for Amyloid Fibril Detection. Methods Appl Fluoresc 2016, 4 (3), 034010. 10.1088/2050-6120/4/3/034010.

(41) Charnley, M.; Gilbert, J.; Jones, O. G.; Reynolds, N. P. Characterization of Amyloid Fibril Networks by Atomic Force Microscopy. Bio Protoc 2018, 8 (4), e2732. 10.21769/BioProtoc.2732.

(42) Sung, J. J.; Pardeshi, N. N.; Mulder, A. M.; Mulligan, S. K.; Quispe, J.; On, K.; Carragher, B.; Potter, C. S.; Carpenter, J. F.; Schneemann, A. Transmission Electron Microscopy as an Orthogonal Method to Characterize Protein Aggregates. J Pharm Sci 2015, 104 (2), 750–759. 10.1002/jps.24157.

